# Systematic Background Selection for Enhanced Contrastive Dimension Reduction

**DOI:** 10.1101/2025.05.12.653541

**Authors:** Kwangmoon Park, Zhongxuan Sun, Ruiqi Liao, Emery H. Bresnick, Sündüz Keleş

**Author notes:** Corresponding author **Correspondence** Sündüz Keleş.

## Abstract

Contrastive dimension reduction enhances the analysis of high-dimensional data by generating target-specific low-dimensional representations relative to a background. Emerging methods for contrastive dimension reduction have demonstrated their utility in extracting target-specific signals across domains involving high-dimensional observations, including genomics, transcriptomics, and pattern recognition. However, even though choosing an appropriate background is critical to the success of contrastive dimension reduction, no established criterion currently exists for selecting such backgrounds. To address this gap, we introduce BasCoD, a novel testing framework grounded in the spectral theory of subspace inclusion, to enable rigorous evaluation and optimal selection of backgrounds. Extensive application of BasCoD across diverse single-cell datasets demonstrates its effectiveness in systematically identifying suitable backgrounds, thereby significantly improving the contrast and interpretability of the derived target representations. We further illustrate how BasCoD can facilitate design of appropriate backgrounds in large-scale single cell experiments under heterogeneous conditions.

## Main

Advances in high-throughput sequencing yield massive genomic datasets across diverse conditions (e.g., treatment *vs*. control). For example, population-scale scRNA-seq^1, 2^ or Perturb-seq^3^ experiments provide high dimensional features that can be compared across different treatments or perturbations. However, their ultra-high dimensionality requires effective dimension reduction to uncover biological insights. In addition, to better contrast high-dimensional datasets from different conditions, a systematic approach is needed to extract low-dimensional features that are specific to the target dataset in comparison to background datasets.

Contrastive Dimension Reduction (CDR)^4–9^ isolates signals specific to a target dataset (e.g., treatment) by comparing them to those of a background dataset (e.g., control) in the low-dimensional space. These CDR methods have demonstrated their effectiveness in deriving target dataset-specific low-dimensional features across a wide range of application areas involving high-dimensional observations, such as genomics, proteomics, and pattern recognition. While numerous CDR methods have been proposed, the selection of an appropriate background remains crucial, as an unsuitable choice may obscure target signals. While these methods typically list suitable examples of background datasets for valid contrastive analysis of a target dataset, a principled criterion to guide this critical selection step has not been established.

To address this critical gap, we introduce BasCoD, which stands for **Ba**ckground **S**election for **Co**ntrastive **D**imension reduction. BasCoD is grounded in the principle that a valid background dataset should not exhibit strong salient structures that are unique or dominant relative to those in the target dataset. This assumption underlies a broad class of contrastive dimension reduction (CDR) methods ^6–9^. Accordingly, BasCoD introduces a formal testing framework to evaluate the suitability of a candidate background dataset by comparing the low-dimensional representations derived from a user-defined dimension reduction method for both the target and background datasets (**Fig**. 1). The resulting decision rule facilitates a statistical goodness-of-fit test for CDR, which empirically aligns with the practical effectiveness of contrastive representations. Beyond its role as a diagnostic tool, BasCoD provides a principled strategy for identifying or designing appropriate background datasets in large-scale, high-dimensional single-cell experiments conducted under heterogeneous conditions. As a result, it supports more reliable downstream analyses by ensuring that contrastive insights reflect meaningful target-specific variation rather than background-driven artifacts. We showcase the effectiveness of BasCoD both on CDR analysis from the literature and with novel applications to single cell trajectory analysis.

**Figure 1.**
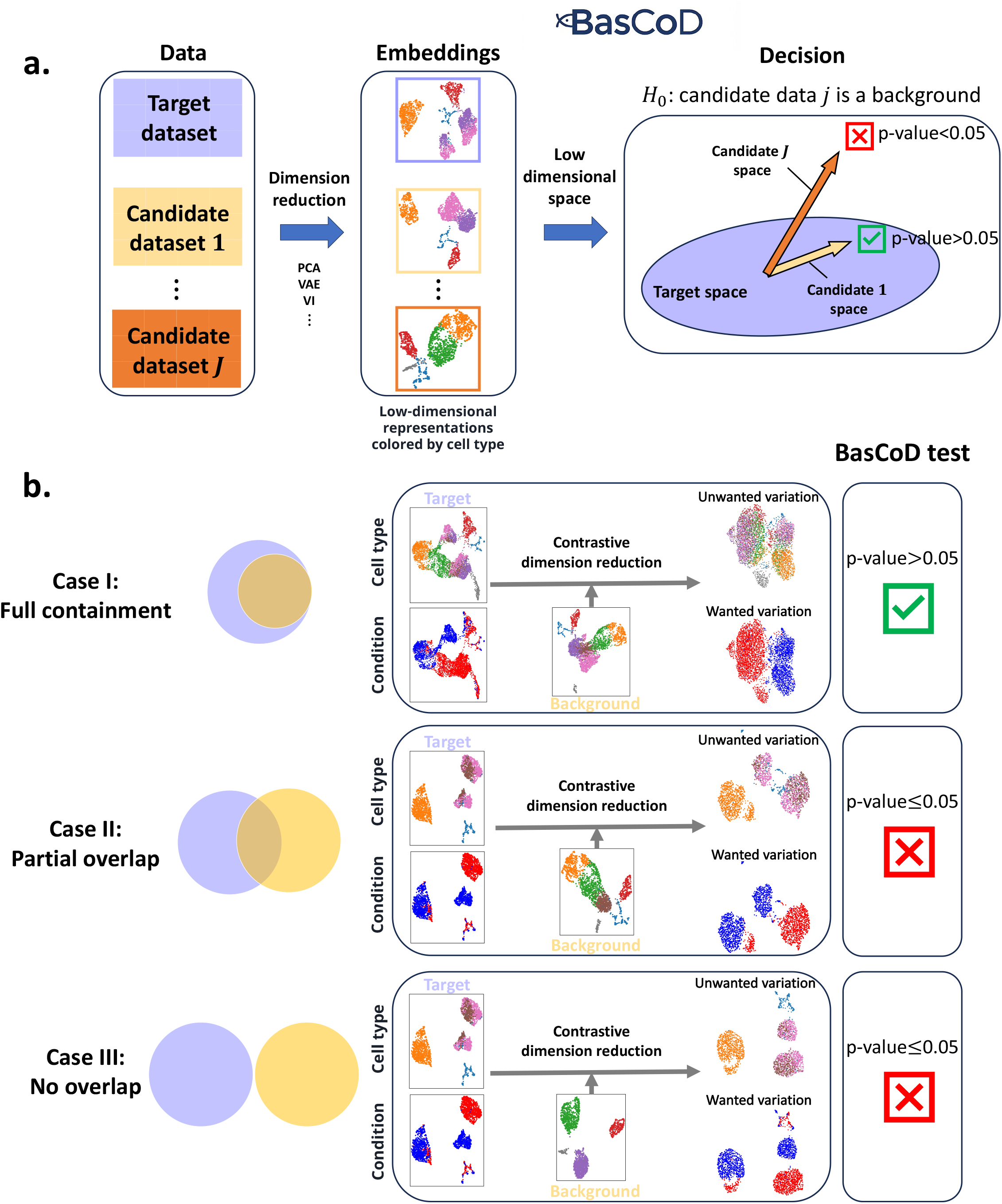
A schematic overview of BasCoD for background dataset selection in contrastive dimension reduction. **a**. A target dataset of interest and a collection of *J* candidate background datasets are considered as inputs (Left). BasCoD utilizes these datasets along with their low-dimensional embeddings derived from a user-selected dimension reduction method (e.g., PCA, VAE, or VI; Middle). It then systematically compares the embedding subspaces of the target dataset to those of each candidate background (Right). Specifically, BasCoD evaluates the null hypothesis *H*_0,*j*_: the embedding subspace of candidate dataset *j* is fully contained within that of the target dataset, for each *j* = 1, …, *J*. **b**. An illustration of typical outcomes when applying BasCoD. Case I (top row) represents an ideal scenario where the candidate background’s embedding subspace (yellow region) is fully included within that of the target (blue region). For this setting, contrastive dimension reduction effectively highlights target-specific variation (e.g., experimental condition) rather than common variation (e.g., cell type), and BasCoD appropriately accepts the null hypothesis (green checkmark). In contrast, Case II and Case III depict scenarios where the candidate backgrounds possess distinct, salient low-dimensional structures differing from the target. In these cases, contrastive dimension reduction is unlikely to enhance target-specific signals, and consequently, BasCoD rejects the null hypothesis (red X mark).

## Results

### The BasCoD testing framework

BasCoD leverages spectral theory of subspace inclusion to assess whether a candidate dataset is a suitable background for contrastive dimension reduction of a target dataset. In practice, when a high-dimensional target dataset is to be contrasted with several candidate background datasets, a dimension reduction method (e.g., Principal Component Analysis (PCA) or Variational Auto Encoder (VAE)) is first applied to each dataset to obtain low-dimensional embeddings (**Fig**. 1a, first row). Then, out of these low-dimensional embeddings, BasCoD derives corresponding low-dimensional spaces (**Methods**) to compare them across datasets (**Fig**. 1a, second row). For effective contrastive analysis, the target must exhibit a more complex low-dimensional structure than the background. We quantify this complexity by determining the dimensionality of the target representation and testing whether the candidate background’s representation lies within the span of the target representation (**Fig**. 1a, decision component). Then, we construct a statistic to assess and formally test this relation based on spectral theories related to subspace inclusion (**Methods**, Algorithm 1). For example, if Candidate background *J* (**Fig**. 1a) forms a distinct salient space, BasCoD returns a small p-value indicating that the candidate is sufficiently different from the target in low-dimensional representation; conversely, if the candidate lacks additional structure, as with Candidate background 1, it yields a large p-value. Specifically, BasCoD rejects the null hypothesis *H*_0_ that a candidate can serve as back-ground when it exhibits extra salient low-dimensional structures (Cases II and III in **Fig**. 1b). Simulation studies demonstrate that the p-values from this testing framework are well calibrated under *H*_0_ (**Fig**. S1a-c, Supplementary Notes) and that detection among non-background datasets maintains controlled false discovery rate (FDR) with high power across varying sub-space perturbation levels which signify the difficulty level of the background selection problem (**Fig**. S1d, Supplementary Notes).

### BasCoD corroborates published contrastive dimension reduction analyses

We first demonstrated BasCoD’s capability to provide a rigorous goodness-of-fit test for results previously reported in the contrastive dimension reduction literature. We examined a mouse protein expression dataset^10, 11^ analyzed in the cPCA study^4^, where mice given shock therapy (target group) were contrasted against control mice (background group). Within the target group, Ts65Dn mice with Down Syndrome relevant neurological phenotypic features (DS) and control (non-DS) mice were nearly indistinguishable using standard PCA (Adjusted Rand Index (ARI) score = 0.01, Normalized Mutual Information (NMI) score = 0.01, average Silhouette score = 0.06, see **Fig**. 2a). However, when the control mice were used as background in a contrastive PCA (cPCA), a clear separation emerged between the DS and non-DS mice (ARI score= 0.78, NMI score = 0.43, average Silhouette score = 0.74, **Fig**. 2a). As emphasized by Abid *et al*.^4^, the control mice capture natural variation in the absence of treatment-induced effects, i.e., a favorable situation essential for a valid background in contrastive dimension reduction. BasCoD directly quantifies the rarity of this favorable situation, and its application to this dataset yields a p-value of 0.16, thereby supporting the validity of the contrastive dimension reduction for the target dataset with the chosen background dataset.

**Figure 2.**
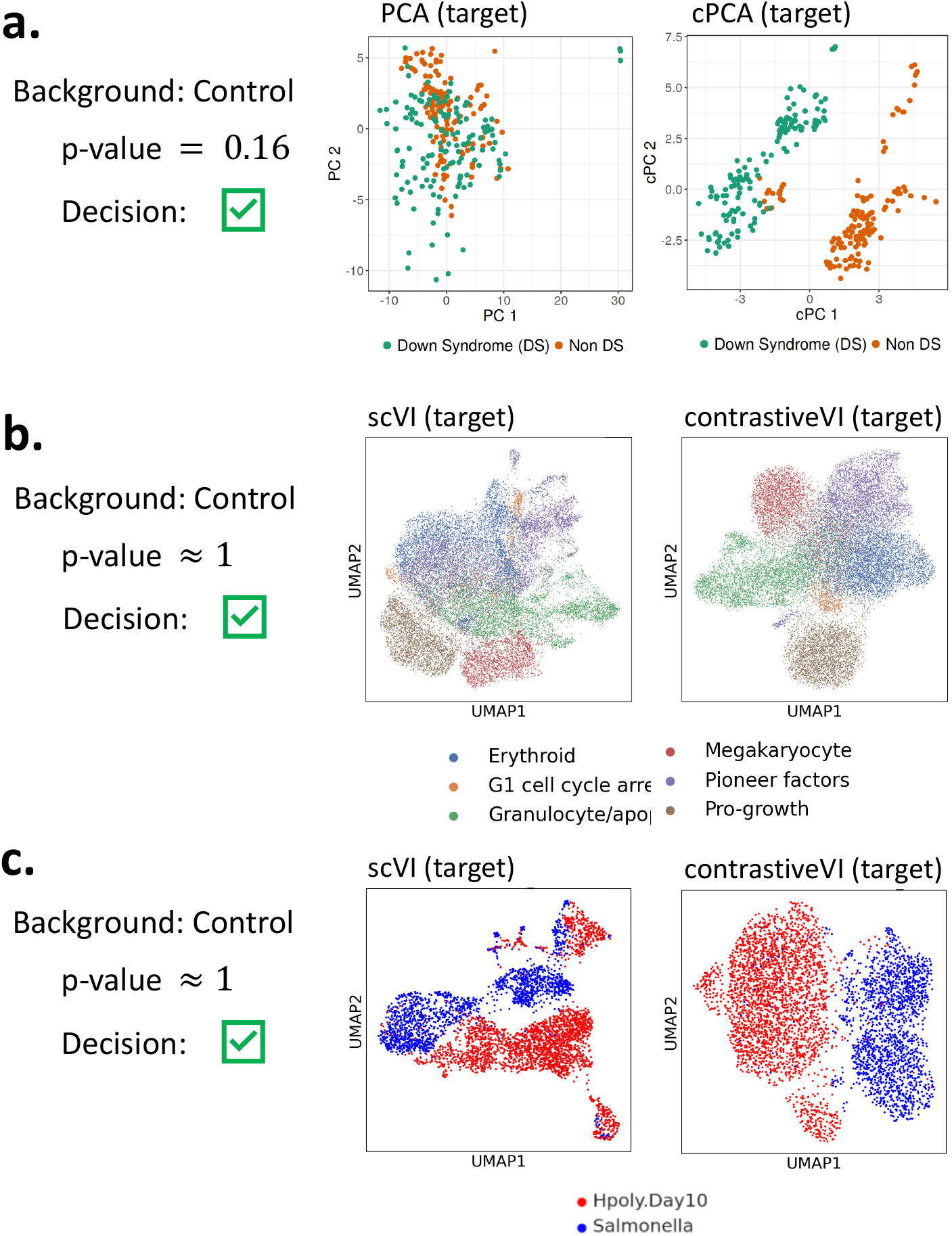
a-c. BasCoD as a goodness-of-fit test for previously reported results from the contrastive dimension reduction literature. **a**. Left: PCA coordinates of the protein expression data of mice that received shock therapy (target dataset) colored by Down Syndrome label. Right: Same figure with cPCA coordinates ^4^ of the target dataset against a group of mice that did not receive shock therapy (background dataset). BasCoD test for the control group mice provides p-value= 0.16, which indicates that the background is valid (green check mark). **b**. Left: UMAP coordinates of the scVI ^13^ cell embeddings obtained from the perturbed cells (target dataset) of ^12^ colored by gene program label. Right: Same figure with contrastiveVI ^7^ salient embeddings, where the target dataset was contrasted against the non-perturbed control cells (background dataset). BasCoD test for the control group mice provides p-value≈ 1. **c**. Left: UMAP coordinates of the scVI embeddings of the entire mouse intestinal cells ^21^ that received pathogen treatments (target dataset). Right: Same figure with contrastiveVI ^7^ salient embeddings, where the target dataset was contrasted against the control cells without pathogen treatment (background dataset).

Having confirmed that BasCoD performs reliably in the linear contrastive dimension reduction setting, we extended its application to a state-of-the-art nonlinear approach, contrastiveVI^7^, to demonstrate its broader applicability. Utilizing a subset of the large-scale Perturb-seq dataset from Norman *et al*.^12^ (filtered as in Weinberger *et al*.), we designated perturbed cells as the target and control cells as the candidate background. Notably, embeddings derived from scVI^13^ exhibited clear associations with cell cycle phase labels in both datasets (**Fig**.S2a), whereas the target dataset additionally revealed subtle separation according to gene-program labels (**Fig**.2b). We subsequently employed BasCoD to rigorously assess whether the low-dimensional loadings of the candidate background, calculated via **eqn**.(4) (**Methods**), were encompassed within the subspace spanned by those of the target data. BasCoD yielded a p-value of ≈ 1, confirming the suitability of control cells as an appropriate background. As expected, contrastiveVI generated salient target-specific embeddings clearly separated according to gene-program labels (**Fig**.2b). We observed similar results in an analysis of a mouse intestinal single-cell dataset originally generated by Haber*et al*.^14^. Here, cells exposed to two pathogens (*H. polygyrus* or *Salmonella*) served as the target, and cells from healthy mice as the candidate background. ContrastiveVI successfully captured pathogen-specific variation in the target dataset, and BasCoD yielded a p-value of ≈ 1, validating the use of healthy cells as an appropriate background (**Fig**. 2c, **Fig**. S3). Taken together, these findings demonstrate that BasCoD corroborates the results of the previous contrastive dimension reduction studies.

### BasCoD offers insights for post-trajectory analysis

Next, we illustrate how BasCoD can yield insights for post-trajectory analysis. Trajectory analysis traditionally involves constructing a minimum spanning tree on a low-dimensional representation of high-dimensional data ^15–17^. Following this, one might wish to contrast the low-dimensional space at a specific time point with that at a branch point of the tree. However, transitions along pseudo-time do not guarantee that data at one time point are optimally contrasted against those at another. In this setting, BasCoD can be leveraged to compare datasets at different time points and identify which pairs are best suited for contrastive analysis. As an illustration, we applied BasCoD to a subset of the Human Cell Atlas bone marrow (HCA-BM) dataset from Hou *et al*., 2023 ^18^. Using TSCAN ^16^, we inferred three hematopoietic stem cell (HSC) lineages - Lymphoid, Myeloid, and Erythroid (**Fig**. 3a) - consistent with Hou *et al*. Firstly, an exploratory analysis based on PCA revealed that HSC cells exhibit higher rank than their differentiated progeny (**Fig**. S4b), suggesting that the lineages have reduced gene expression variation relative to HSC. We thus designated HSC as the target dataset and treated each lineage cluster as a candidate background. Applying BasCoD to each cluster, we found that only the two clusters corresponding to the Erythroid lineage (green and yellow in **Fig**. 3a) were unsuitable as backgrounds. Specifically, BasCoD yielded p-values of 0.29 for the Lymphoid lineage, 0.6 for the Myeloid lineage, and 0.005 and 0.0004 for the two Erythroid clusters, indicating that the Erythroid lineage exhibits the most drastic deviation in low-dimensional gene expression structure from HSC. To support these findings biologically, we examined the expression patterns of the top genes, SRSF7 and TPM4, which contribute to the first two principal components of the HSC data. In the Erythroid lineage, these genes showed significant differences in expression relative to HSC compared to the other lineages (**Fig**. 3b). This suggests that lineage-specific variations in the Erythroid branch may confound contrastive dimension reduction, underscoring the importance of appropriate background selection.

**Figure 3.**
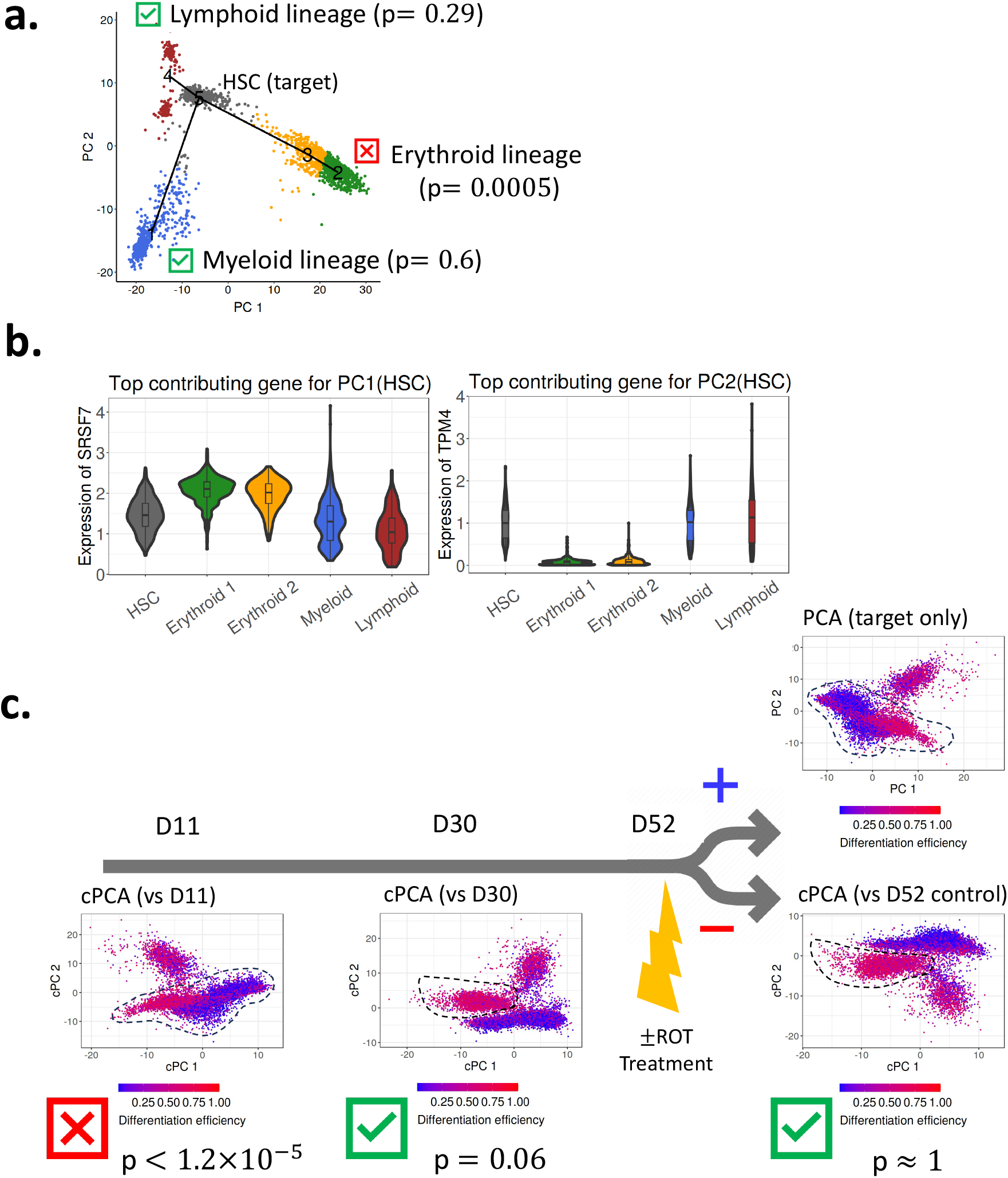
Background selection for post-trajectory analysis contrastive dimension reduction with BasCoD. **a**. PCA coordinates of a subset of HCA-BM cells ^18^ with the TSCAN ^16^ minimum spanning tree. Each lineage (candidate background dataset) learned from TSCAN is evaluated by BasCoD for a contrastive analysis against the HSC cells (target dataset) and BasCoD p-values are reported for each lineage. **b**. Expression levels of the top gene with the largest weight for PC1 (or PC2) across cell types of ^18^ dataset. **c**. Top right: PCA coordinates of the human iPSCs at D52 with ROT treatment (target dataset), colored by differential efficiency level of each donor. Bottom right: cPCA coordinates of the same D52 (ROT) cells after cPCA against control cells at D52 (candidate background dataset). Cells marked with dashed-line circle represent the ones from donors with high differentiation efficiency. methodname p-value is reported at the bottom of the PCA plot. Middle and left panels depict the same plots with D30 and D11 cells as candidate background datasets, respectively.

To further illustrate the pitfalls of invalid background selection in contrastive dimension reduction along time trajectories, we analyzed population-scale scRNA-seq data of human induced Pluripotent Stem Cells (iPSCs) differentiating toward a midbrain neural fate measured at Day 11 (D11), Day 30 (D30), and Day 52 (D52) ^1^. At D52, a subset of cells was exposed to rotenone-induced oxidative stress (ROT), while the remaining cells served as controls (**Fig**. 3c). Furthermore, for each of the 213 donor iPSCs, Jerber *et al*. defined a *Differentiation Efficiency (DE)* value, representing the proportion of dopaminergic and serotonergic-like neurons at D52, to capture donor-specific differentiation variation. We defined ROT-treated cells at D52 as the target dataset and considered three candidate backgrounds: cells at D11, D30, and D52 (control). When PCA was applied to the target dataset alone, cells from high and low DE donors were intermingled (dash-lined cells in the top right panel of **Fig**. 3c, *R*^2^ = 0.26 in the regression of DE on the two cPCs). However, cPCA using D30 and D52 (control) as backgrounds revealed distinct clusters separating high and low DE donors with *R*^2^ = 0.35 and 0.36, respectively (middle and right panels of **Fig**. 3c). In contrast, using D11 cells as the background did not yield such separation with *R*^2^ = 0.28 (leftmost panel of **Fig**. 3c). Evaluating background validity with BasCoD further supported these findings. The p-values for D11, D30, and D52 (control) were 1.2 × 10^−5^, 0.06, and ≈ 1, respectively, indicating that D30 and D52 (control) are valid backgrounds for the D52 ROT-treated cells. This is consistent with Jerber *et al*.’s observation that cell type proportions at D11 differ dramatically from those at later time points, whereas D30 and D52 cells are relatively homogeneous. In summary, these results demonstrate that BasCoD provides a critical tool for verifying the reliability of contrastive analyses on low-dimensional features across time points, whether defined by experimental design or inferred from trajectory analyses.

### BasCoD enables background data calibration in single-cell experiments

After demonstrating BasCoD’s ability to evaluate whether a given dataset can serve as a suitable background for contrastive dimension reduction of a target dataset, we now illustrate how it can further guide the design and refinement of candidate backgrounds. As detailed in the **Methods** section, a valid background according to BasCoD should not harbor any distinct low-dimensional features beyond those present in the target dataset, and its low-dimensional embedding space must lie entirely within that of the target (**Fig**. 1a). To exemplify a pathological scenario, we revisited the mouse intestinal single-cell dataset, constructing target and candidate datasets with completely non-overlapping cell types (**Fig**. 4a). Under these conditions, contrastiveVI fails to reveal any additional insights on pathogen types, as its salient embeddings show (**Fig**. 4b). Correspondingly, BasCoD returned an extremely low p-value (3.2 × 10^−200^), clearly rejecting the candidate background. As an intermediate scenario, we constructed a candidate background dataset containing both cell types common to the target (Endocrine, Enterocyte, and TA) and candidate-specific cell types (Goblet, Enterocyte.Progenitor, and Tuft), highlighted by black dashed circles in **Fig**. 4c. Here, the target dataset also exhibited distinct variation within TA cells (red dashed circles in **Fig**. 4c). Although contrastiveVI improved pathogen-type clustering compared to the target-only dimension reduction (ARI score = 0.2, NMI score = 0.18, average Silhouette score = 0.47), the clustering was still suboptimal. Consistently, BasCoD yielded a p-value of 3.6 × 10^−11^, indicating the candidate background was not valid.

**Figure 4.**
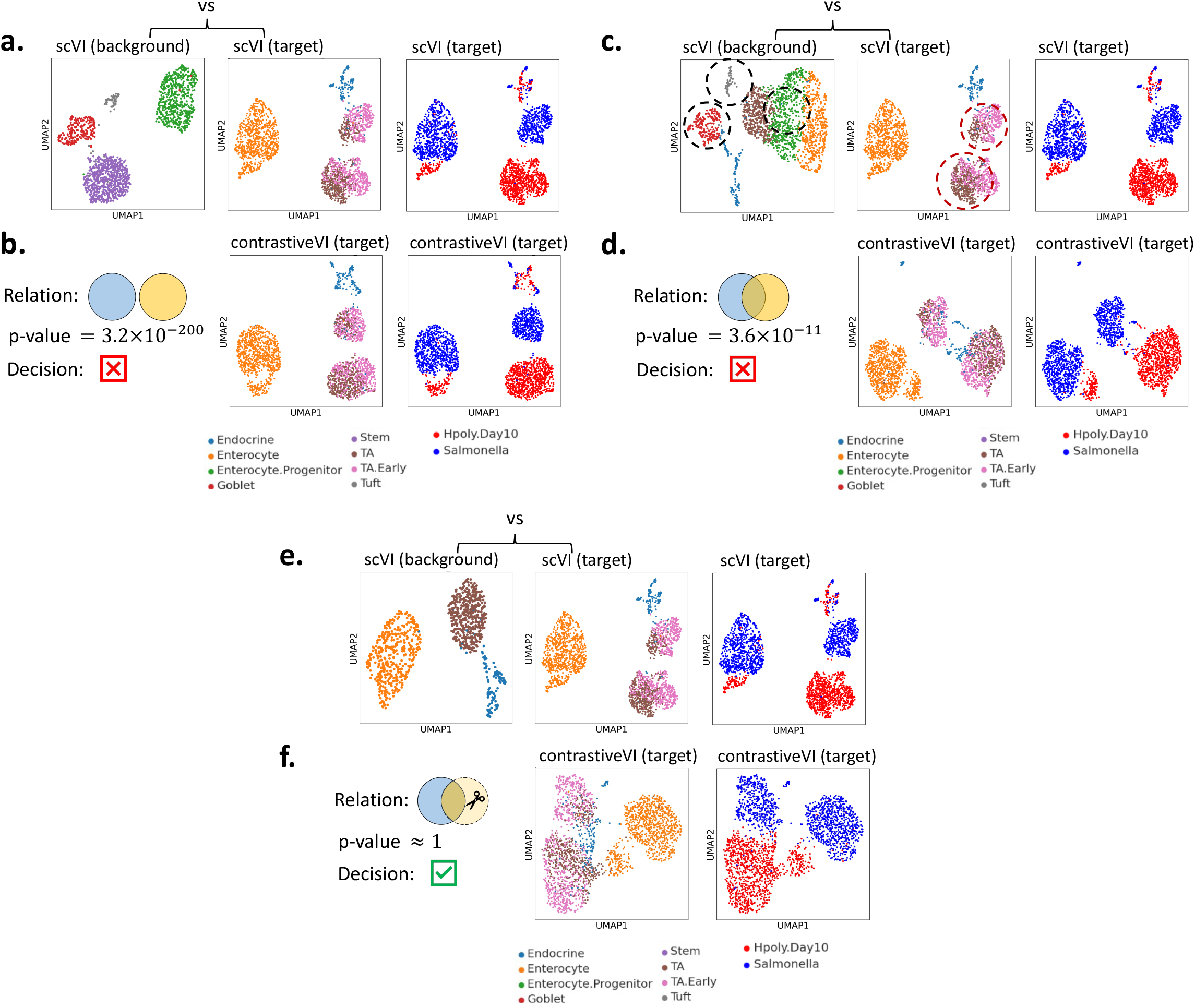
Background data calibration for improved contrastive dimension reduction with BasCoD. **a**. Left: UMAP coordinates of the scVI embeddings of a subset of mouse intestinal cells ^21^ without pathogen treatment (candidate background designed to have no overlapping cell types with the target dataset) colored by cell type. Middle: same figure with the cells that received pathogen treatments (target dataset). Right: Same figure as the middle panel, but colored by the two of the pathogen types. **b**. Left: Venn-diagram depicts the designed relation between target and candidate background. A BasCoD p-value with a decision on the candidate background is reported. Middle: UMAP coordinates of contrastiveVI salient embedding of the target dataset contrasted against the candidate background dataset, colored by cell type. Right: same as the middle panel, but colored by pathogen type. **c**. Left: UMAP coordinates of the scVI embeddings of a subset of mouse intestinal cells ^21^ without pathogen treatment (candidate background) colored by cell type. The cells within dashed-line circles are the ones that are only present in the candidate background dataset. Middle: same figure with the cells that received pathogen treatments (target dataset). Right: Same figure as the middle panel, but colored by the two of the pathogen types. **d**. Left: Venn-diagram depicts the designed relation between target and candidate background. A BasCoD p-value with a decision on the candidate background is reported. Middle: UMAP coordinates of contrastiveVI salient embedding of the target dataset contrasted against the candidate background dataset, colored by cell type. Right: same as the middle panel, but colored by pathogen type. **e**,**f**. Same figures as **c**,**d**., but with calibrated candidate background dataset by removing the cells (circled cells in **e**) that are specific only to the candidate background dataset.

Following this rejection, we applied a sequential refinement procedure by removing candidate-specific cell types and reassessing background validity using BasCoD. Specifically, we recalibrated the candidate dataset by excluding the Goblet, Enterocyte.Progenitor, and Tuft cell types (black dashed circles in **Fig**. 4c), separately applied scVI to the target and calibrated candidate datasets, and carried out testing with BasCoD (**Fig**. 4e). This recalibrated candidate yielded a p-value of ≈ 1, affirming its suitability for contrastive analysis. Correspondingly, the contrastiveVI salient embeddings derived from this refined background exhibited markedly improved separation by pathogen type (ARI score = 0.30, NMI score = 0.34, average Silhouette score = 0.45) compared to embeddings obtained from the target-only scVI (**Fig**. 4f). In summary, these results demonstrate that BasCoD not only rigorously assesses candidate backgrounds but also provides a systematic framework for their effective design and calibration in contrastive dimension reduction.

### Practical utility of BasCoD for designed experiments with high-dimensional data

Finally, we demonstrate the practical utility of BasCoD for designing valid contrasts in single-cell experiments. In an erythroid differentiation analysis with primary human hematopoietic stem/progenitor cells (HSPCs), cells were subjected to culture under conditions or upon inflammation with a mixture of the cytokines TNF*α*, IFN*γ*, and IL-6 on day 4 of the culture and harvested for single-cell RNA sequencing on days 9 (D9) and 13 (D13). An initial exploratory analysis revealed that, while control cells differentiate as a function of time, inflammation largely blocks differentiation (**Fig**.S5). We focused on cells that differentiated by D13 (orange cells within the dashed circle in **Fig**.S5), defining them as “escaped cells” post-inflammation. We set these D13 inflammation-treated cells as the target and considered three candidate backgrounds: inflammation D9, control D9, and control D13. PCA on the target dataset alone failed to separate the escaped cells (ARI = 0.18, **Fig**.S6a), a misalignment also evident with the Leiden clustering of the principal components (**Fig**.S6b, **Fig**.S7). We applied cPCA^4^ to contrast the target against each candidate background. When contrasting inflammation-treated D13 and D9 cells (Contrast I in **Fig**. 5a), the escaped cells were clearly separated (ARI = 0.69), and BasCoD returned a p-value of ≈ 1, confirming that inflammation-treated D9 cells represents a valid background. This supports our expectation that fixing one factor (treatment) while comparing different times enhances the capacity to identify temporal effects. In contrast, comparing inflammation-treated D13 cells with control D13 cells (Contrast III in **Fig**. 5a) yielded poor separation (ARI = 0.16) and a BasCoD p-value of *<* 2 × 10^−146^, due to the candidate’s strong, independent low-dimensional features (as evident by the sparse distribution of earlier stage cells in **Fig**.S5). Similarly, using inflammation-treated D9 cells as the candidate (Contrast II in **Fig**. 5a) resulted in a p-value *<* 0.001 and only moderate clustering (ARI = 0.49). Notably, the UMAP embedding of inflammation-treated D9 cells is similar to that of control D9 cells, but with fewer late-stage cells, reinforcing its validity as a background. Collectively, this analysis highlights the capacity of BasCoD to identify and calibrate appropriate background dataset, thereby avoiding misleading inferences in contrastive dimension reduction. Critically, inappropriate background selection can lead to misleading downstream findings. For instance, Gene Set Enrichment Analysis (GSEA)^19^ on cell clusters derived from cPCA with a valid background (D9 with inflammation) showed that escaped cells exhibit significantly lower expression of immune function genes (e.g., leukocyte and lymphocyte activation) in comparison to other cells (**Fig**. 5b). This result was consistent with a gold standard GSEA based on inferred cell labels, with seven out of ten gene sets matching (**Fig**. 5c). In contrast, GSEA on other contrasts matched only one out of ten gene sets (**Fig**. S8a,b), emphasizing that invalid background selection can lead to incomplete/misleading inference.

**Figure 5.**
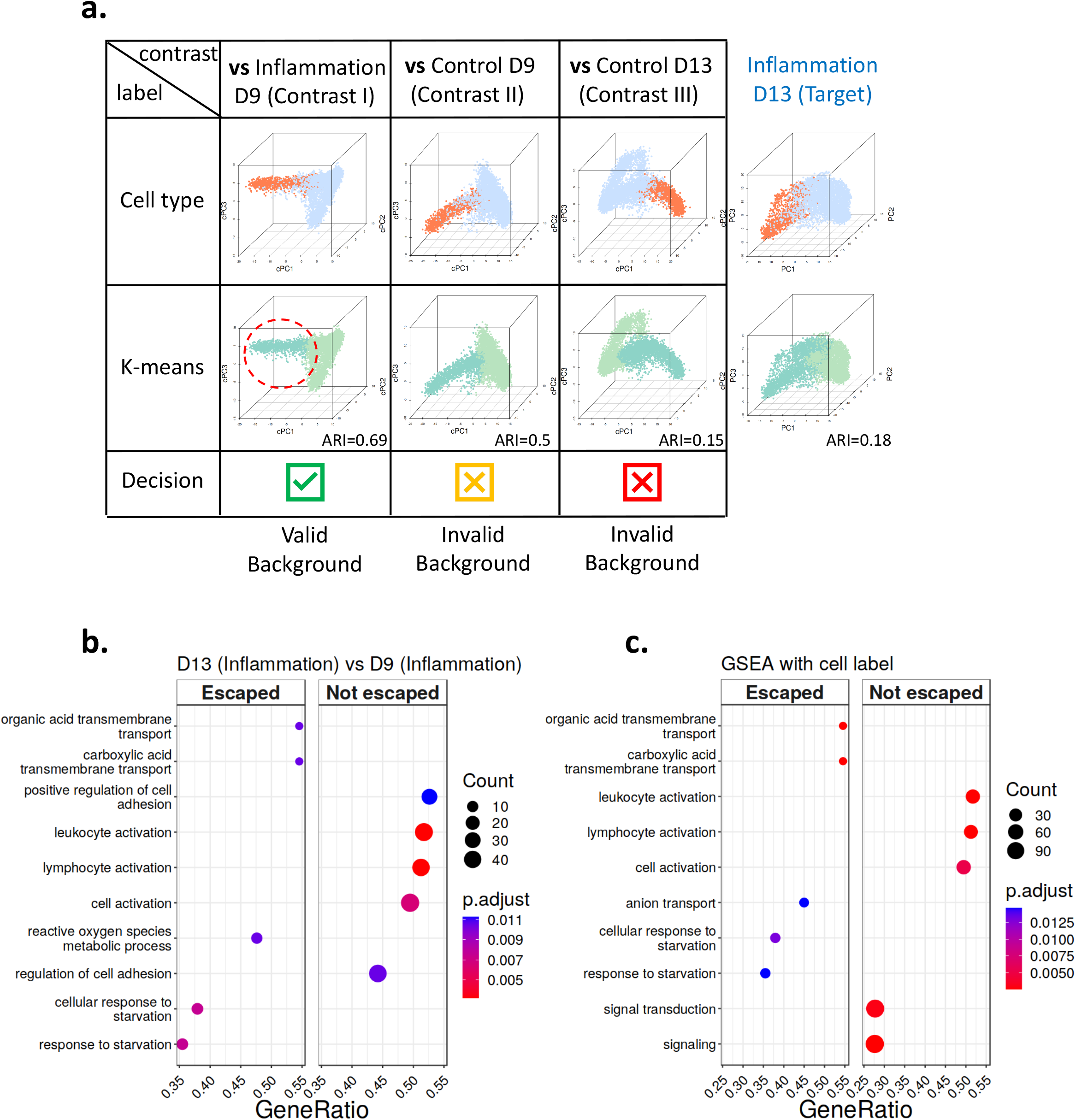
Application of BasCoD for background selection in a designed scRNA-seq experiment. **a**. The right-most column depicts PCA embeddings (first three components) of HSPC cells at day 13 (D13) under inflammation treatment (target dataset), colored by cell types (top) and k-means derived cell clusters (bottom). The ARI at the bottom quantifies the concordance between cell type labels and clustering. Each column in the table depicts cPCA embeddings contrasting the target dataset against candidate backgrounds: D9 cells with inflammation, D9 cells without inflammation, and D13 cells with inflammation. The BasCoD decisions are reported in the bottom row. **b**. Gene set enrichment analysis (GSEA) comparing the two cell clusters derived from contrastive PCA embeddings when the target dataset (D13 with inflammation) is contrasted against inflammation-treated D9 cells. The “Escaped” panel contains the gene functions enriched in escaped cells and the “Not escaped” panel contains those enriched in not escaping cells. **c**. GSEA result comparing escaped and not escaping cells of the target dataset (D13 with inflammation).

## Discussion

We introduced BasCoD which successfully exploits the low-dimensional spaces of target and candidate background datasets to provide a goodness-of-fit test for a contrastive dimension reduction with large-scale heterogeneous genomic datasets. The p-values from the BasCoD testing framework are well calibrated under the null hypothesis that a dataset is a valid background. In addition, BasCoD exhibit well controlled FDR and high power in detecting invalid background datasets from sets of simulation experiments. These results signify the reliability of the BasCoD testing framework. A recently proposed method ^20^ specifically evaluates the presence of additional low-dimensional structures in the target dataset relative to a candidate background. Although this approach effectively identifies whether the target dataset contains structure distinct from the the background, it does not address the crucial task of validating the suitability of the candidate background itself. Importantly, this method is not designed to reject candidate backgrounds whose low-dimensional structures differ entirely from those of the target dataset, as exemplified by the pathological scenario depicted in **Fig**. 4a (Supplementary Notes). For determining the dimensionality of the target and candidate background embeddings, we relied on empirical common practices with exploratory analysis (Supplementary Notes). However, a more systematic way of estimating the dimension would be beneficial. In addition, while we relied on an approximation based approach to derive subspaces of each dataset for non-linear CDR, a non-parametric approach could provide further generalization. For example, comparing the reconstruction loss using the encoder/decoder learned from the candidate background dataset to that using the encoder/decoder learned from the target dataset, through a non-parametric test, could provide insights into subspace inclusion relationship. BasCoD’s results not only corroborate the previous findings from contrastive dimension reduction literature but also facilitate contrastive analysis on learned pseudo-time after a trajectory analysis and the design of background datasets. Moreover, BasCoD is compatible with any dimension reduction method, positioning it as a pivotal tool for advancing contrastive analysis.

## Competing Interests

The authors declare no competing financial interests.

## Methods

### Latent variable models of target and background datasets

Throughout the paper, we denote the *target dataset* as 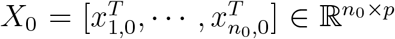, where 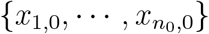 is *n*_0_ i.i.d. copies of *x*_0_ ∈ ℝ^*p*^, and candidate background dataset *j, j* ∈ 𝒞, as 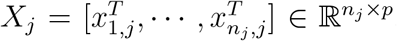, where 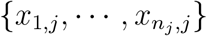 is *n*_*j*_ i.i.d. copies of *x*_*j*_ ∈ ℝ^*p*^. Here, 𝒞 is the index set for the candidate datasets. We model *x*_*j*_ as a function of two sets of latent variables named as (i) data-common embeddings 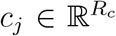 and (ii) data-specific embeddings 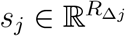, and a random noise *ϵ*_*j*_ ∈ ℝ^*p*^, for each *j* ∈ {0} ∪ 𝒞 as:

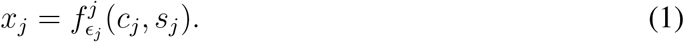

In the contrastive dimension reduction framework, we expect the target dataset to be generated from both *c*_0_ and *s*_0_, and an appropriate background dataset to be generated only through *c*_*j*_. Based on this notion, we define an appropriate *background dataset* as a candidate dataset satisfying *s*_*j*_ = 0, i.e.,

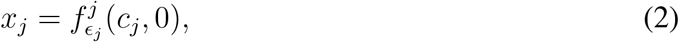

and denote the data index set ℬ := {*j* ∈ 𝒞|*s*_*j*_ = 0}. Then, the main task is to identify the set ℬ ⊂ 𝒞, utilizing the low-dimensional embeddings learned from *X*_*j*_ for *j* ∈ {0} ∪ 𝒞.

It is worth noting that model (1) with a single background dataset, i.e., ℬ = {1}, encompasses a large class of contrastive dimension reduction method models. For example, when 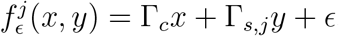, with orthogonal [Γ_*c*_, Γ_*s,j*_], we have

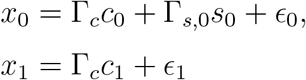

as target and background datasets, respectively. This corresponds to Contrastive Latent Variable Model (CVLM)^6^ and a specific case of Contrastive PCA (cPCA)^4^ model. Here, the loading matrix Γ_*s*_ encodes the low-dimensional space specific to the target dataset and Γ_*c*_ encodes the low-dimensional space common to both of the datasets. When 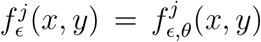 is constructed through a Variational Auto Encoder or Variational Inference with unknown parameters *θ*, the model corresponds to that of ContrastiveVAE^5^ or ContrastiveVI^7^.

### Statistical test for selecting background datasets

With the set up outlined above, we now introduce the procedure of selecting ℬ out of 𝒞. We first illustrate the testing procedure for linear 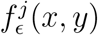 and describe how a similar procedure can be implemented in general. Consider a linear latent variable model 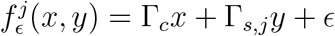 with 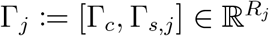. Then, we have the datasets

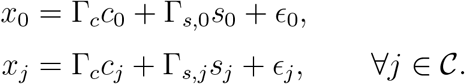

When the *j*th candidate dataset is a valid background, i.e., *j* ∈ ℬ, we have

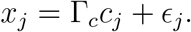

This implies that the candidate’s subspace Γ_*j*_, *j* ∈ ℬ, must match or be contained in the same subspace that the target dataset occupies (Γ_*c*_): *C*(Γ_*j*_) = *C*(Γ_*c*_) ⊂ *C*(Γ_0_), as Γ_*j*_ = Γ_*c*_ for *j* ∈ ℬ, which equivalently implies Γ_*j*_ = *P*_0_Γ_*j*_, where 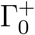 denotes a Moore-Penrose inverse of Γ_0_ and *P*_0_ is the projection matrix onto the space *C*(Γ_0_). If this condition fails, the candidate background dataset *j* introduces additional, unwanted variation and thus is not a suitable background. As a result, we formulate the null hypothesis 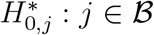 as *H*_0,*j*_ : Γ_*j*_ = *P*_0_Γ_*j*_, which holds if and only if

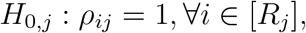

where *R*_*j*_ = *R*_*c*_ + *R*_Δ*j*_, *ρ*_*ij*_ = corr(*γ*_*ij*_, *P*_0_*γ*_*ij*_), corr(·, ·) denotes the sample correlation, and *γ*_*ij*_ denotes the *i*th column vector of Γ_*j*_.

Testing whether the *j*th candidate is an appropriate background requires dimension reduction for each *j* ∈ {0} ∪ 𝒞 to obtain 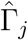 and evaluation of how close 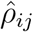 is to 1 for ∀*i* ∈ *R*_*j*_, where 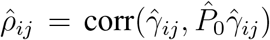. For such evaluation, we consider the Fisher transformation on each correlation estimate 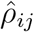 to generate a z-score 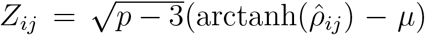, where *µ* = arctanh(1 − *ϵ*) for a small *ϵ >*0 (to avoid *µ* = ∞). Then, under the null hypothesis *H*_0,*j*_, we have each *Z*_*ij*_ ∼ *N* (0, 1) for all *i* ∈ [*R*_*j*_]. Leveraging Fisher’s method for combining p-values, we have

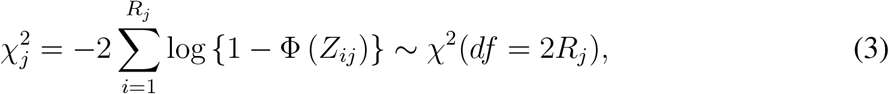

where *df* denotes degrees of freedom and Φ(·) denotes the standard normal cumulative distribution function. While the Fisher’s method to combine p-values rely on independence of the individual p-values across *i* ∈ [*R*_*j*_], variants of this procedure for accommodating dependencies among p-values are available ^14, 22^. With this null distribution for the chi-squared statistic of the combined p-values, 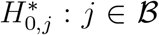 (candidate dataset *j* is a background) is rejected as an appropriate background for small p-values.

When 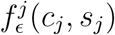 is in a general form (i.e., as in contrastiveVI), a direct analogue of Γ_*j*_ may not be available unless *f*^*j*^ contains a partially linear format depending onthe latent embeddings *c*_*j*_ and *s*_*j*_. As a result, to adopt the BasCoD testing procedure for these settings, we estimate Γ_*j*_ through the following linear approximation:

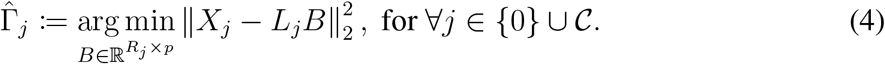

where 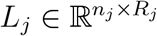 is any low-dimensional embedding obtained from data *X*_*j*_. For contrastive PCA, *L*_*j*_ corresponds to the *R*_*j*_ principal components of *X*_*j*_ and for contrastiveVI or VAE, *L*_*j*_ correspond to *R*_*j*_ latent embedding variables using only *j*th dataset. Then, using 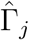 across *j* ∈ {0} ∪ 𝒞, we can follow the same procedure introduced above. Algorithm 1 summarizes the entire BasCoD testing procedure. After constructing/selecting a valid background dataset {*X*_*j*_|*j* ∈ ℬ}, one can proceed with contrastive dimension reduction against the target dataset *X*_0_. The testing procedure requires two key parameters: the size *R*_*j*_ of the low-dimensional space of each *j* ∈ {0} ∪ 𝒞 and the slack parameter *ϵ >* 0. We refer to Supplementary Notes for details on how to set these parameters.

It is also instructive to consider the setting where the target subspace is entirely contained within the candidate background subspace, i.e., *C*(Γ_0_) ⊂ *C*(Γ_1_). In this setting, as the contrastive analysis of the target dataset relative to the candidate background is unlikely to yield additional insights, BasCoD is expected to deem the contrastive dimension reduction invalid. As an illustrative example, when Γ_0_ = Γ_1_[, 1], we expect 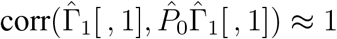, whereas 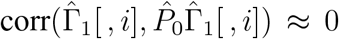 for *i* ≠ 1, leading to the rejection of BasCoD’s null hypothesis. Combined with the settings in **Fig**. 4, this underscores BasCoD’s capacity to assess the validity of contrastive dimension reduction across varied contexts.

#### Algorithm 1 BasCoD

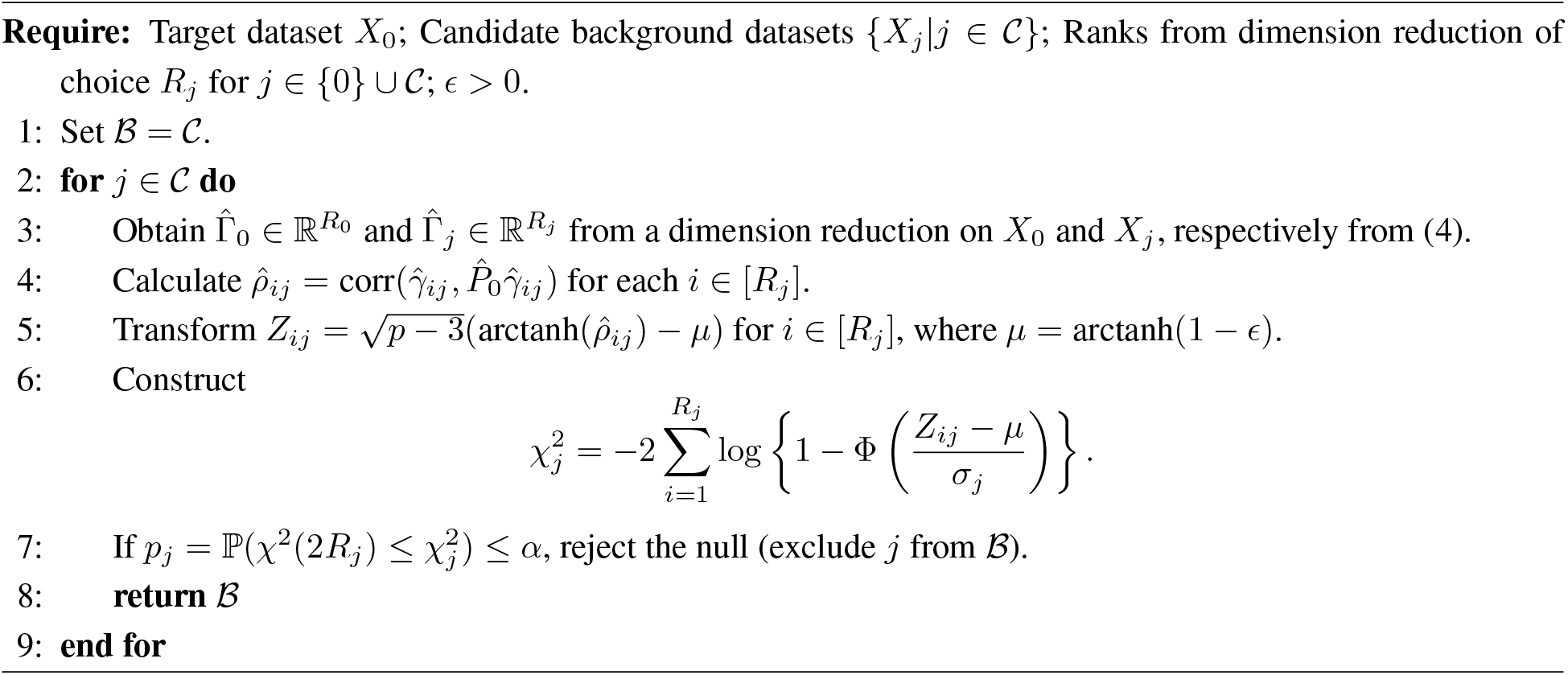

## Code Availability

The BasCoD implementation R code is publicly available on our GithHub (https://github.com/keleslab/BasCoD). The GitHub repository also includes a tutorial that reproduces some of the results reported in the manuscript.

## Data Availability

The datasets used for this manuscript and the sources for them are as follows:

- Mouse protein data^10, 11^: [cPCA^4^ GitHub repository](https://github.com/abidlabs/contrastive/blob/master/experiments/datasets/Data_Cortex_Nuclear.csv).
- Perturb-seq data^12^ : [ContrastiveVI^7^ tutorial](https://figshare.com/ndownloader/files/43390776).
- Mouse intestinal data^21^: [ContrastiveVI^7^ tutorial](https://drive.google.com/uc?id=1-QRE2up3YecF2yrMj1bd_SUQXF_w0RDZ).
- Human Cell Atlas bone marrow (HCA-BM) dataset^18^: [Lamien^18^ tutorial](https://github.com/Winnie09/Lamian/blob/master/data/slingshot_data.rda).
- Population scale single cell RNA-seq data with ROT treatment^1^: [Zenodo link of Jerber *et al*.^1^](https://zenodo.org/records/4333872).
- Single cell RNA-seq dataset with inflammation treatment: Google drive link.

## Supplementary Figures and Tables

**Supplementary Figure 1.**
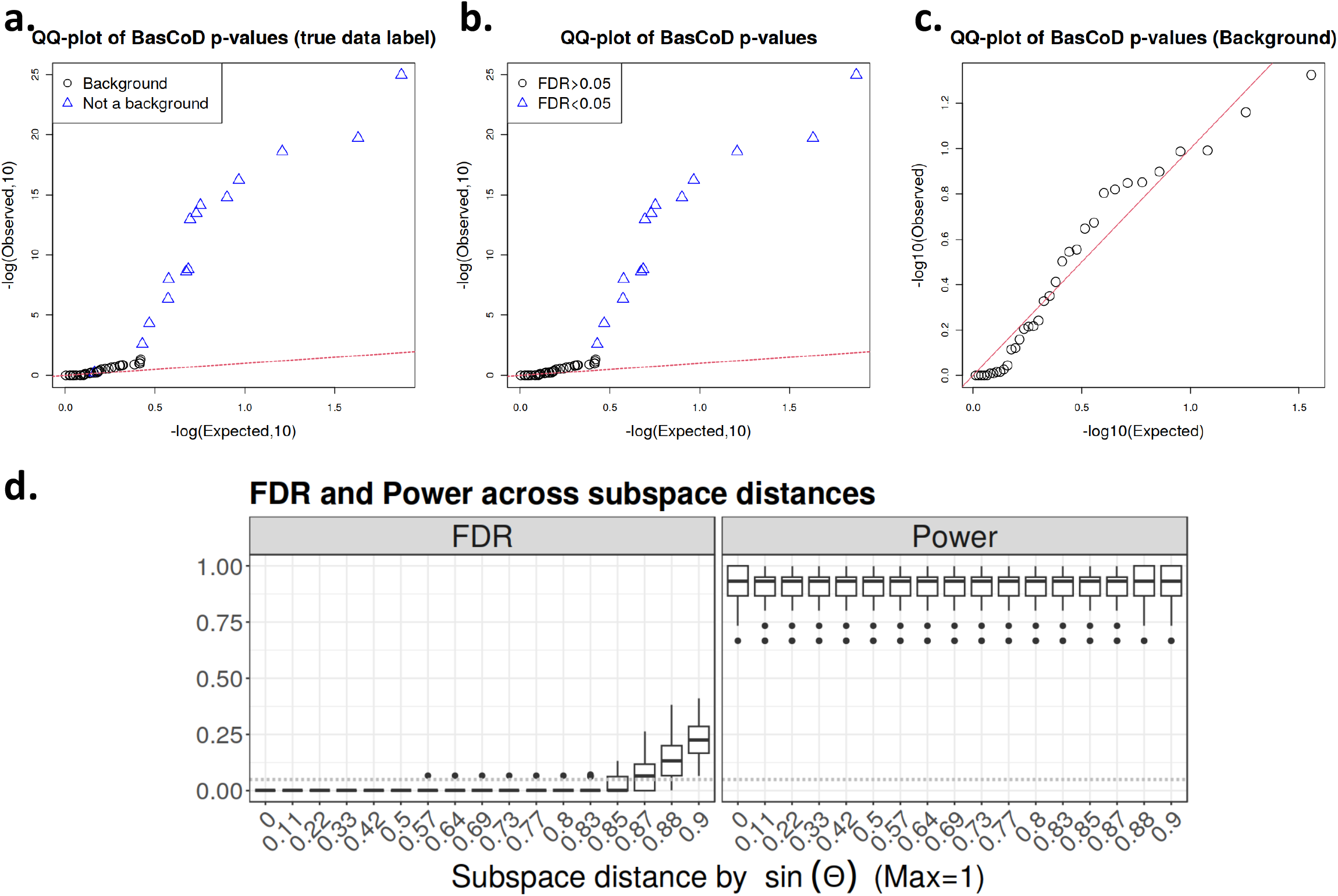
Simulation studies with synthetic datasets. **a, b**. Quantile-Quantile (Q-Q) plots of the BasCoD p-values for 50 candidate background datasets on one simulation example with sin(Θ) = 0.87 compared against the uniformly distributed random variables. Each point represents a candidate background dataset colored by the cell type label (**a**) and the label inferred from BasCoD (**b**). **c**. Same plot for only the 35 background datasets. **d**. Left : empirical False Discovery Rate (FDR) for each sin(Θ) distance perturbation level evaluated across 100 replications. Right: Analogous plot recording power of the test.

**Supplementary Figure 2.**
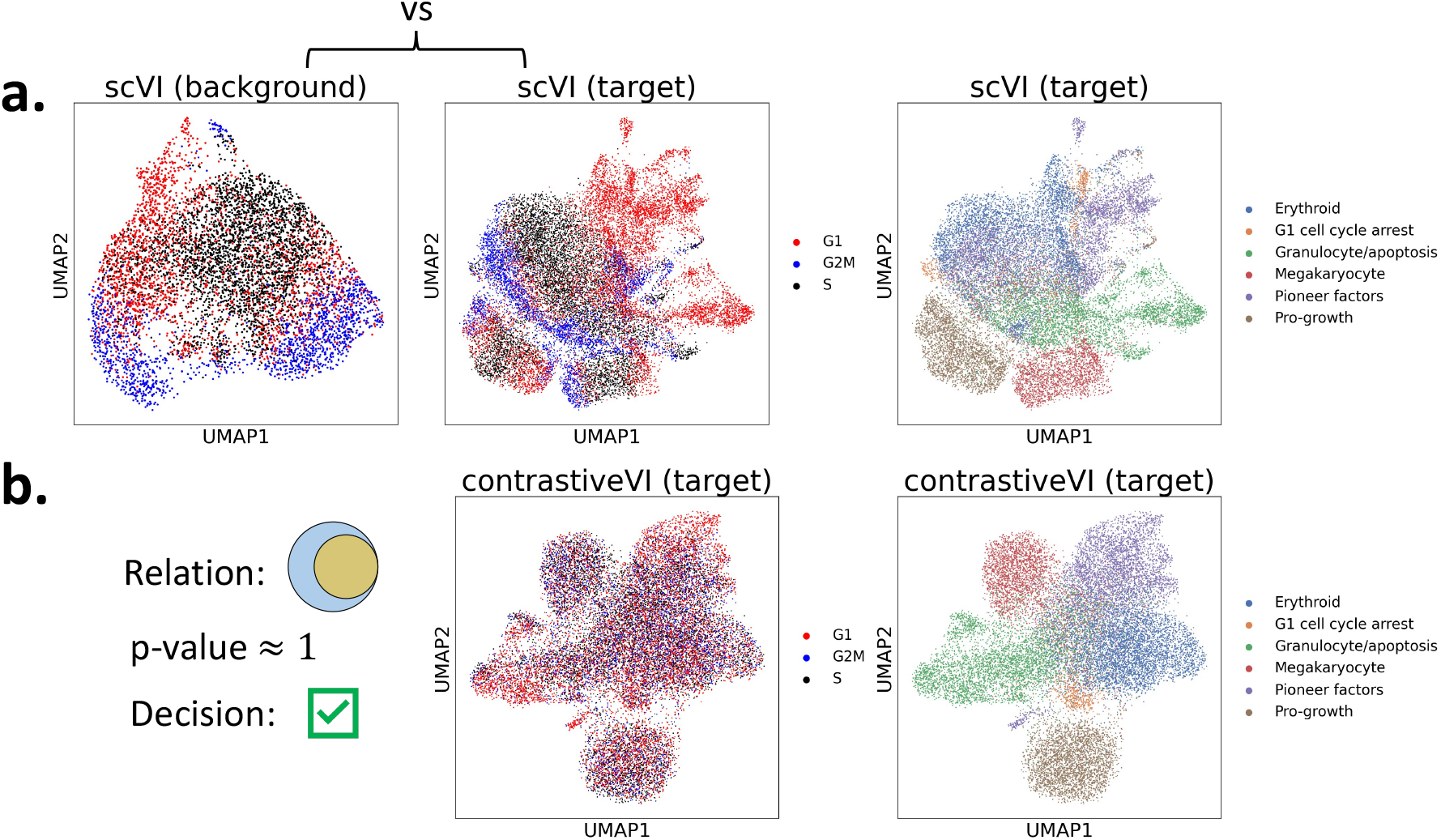
**a**. Left: UMAP coordinates of scVI-derived embeddings for unperturbed cells (candidate background dataset) from the Perturb-seq^12^ experiment, colored by cell cycle phase. Middle: UMAP coordinates of scVI-derived embeddings for perturbed cells (target dataset) from the Perturb-seq^12^ experiment, colored by cell cycle phase. Right: Middle panel embedding colored by the gene program labels. **b**. Left: Venn diagram illustrating the designed relationship between target and candidate background datasets, along with the corresponding BasCoD p-value and the decision regarding the candidate background validity. Middle: UMAP embeddings showing contrastiveVI salient features of the target dataset contrasted against the candidate background, colored by cell cycle phase. Right: Same embedding as the middle panel, colored by the gene programs.

**Supplementary Figure 3.**
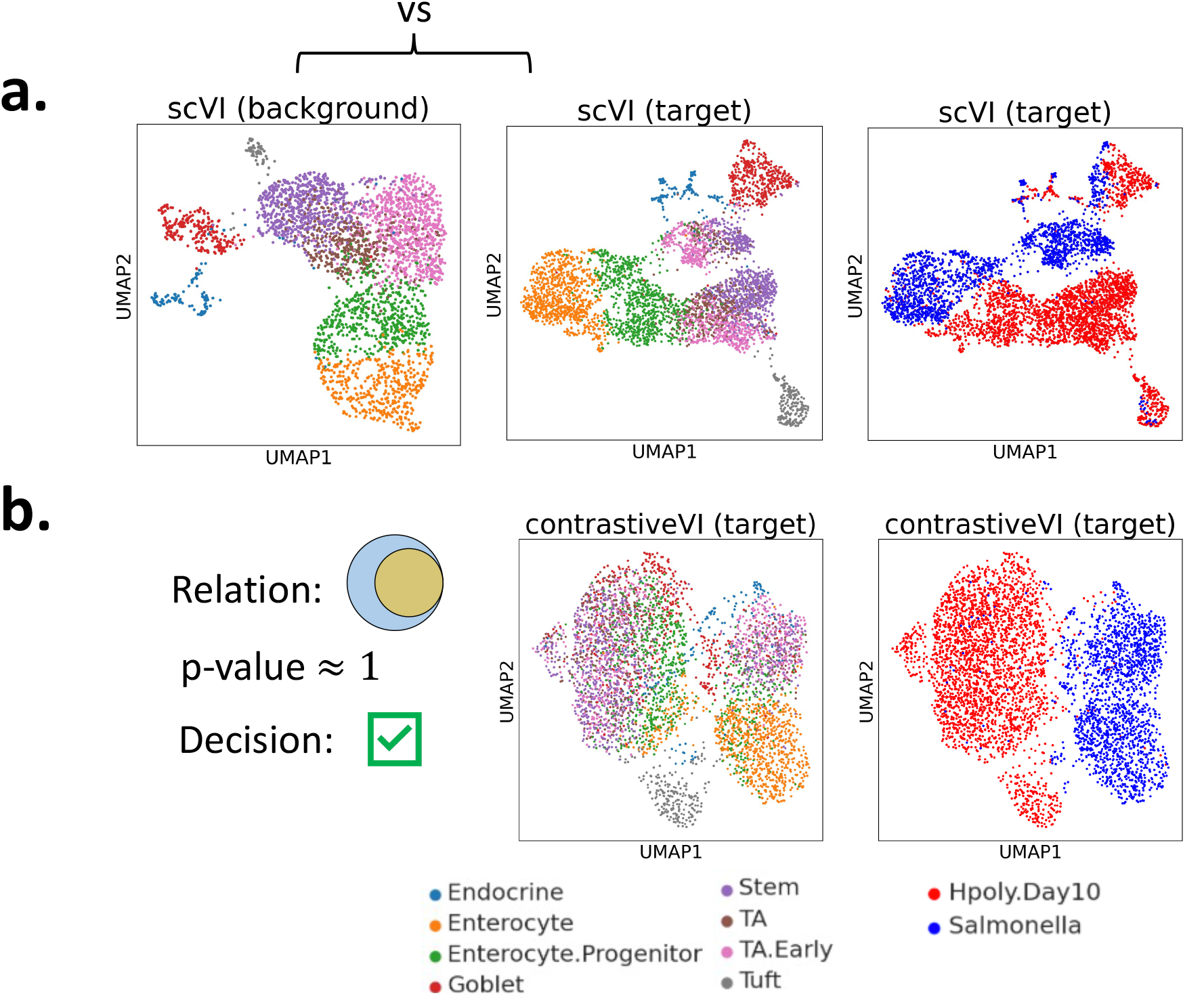
**a**. Left: UMAP coordinates of scVI-derived embeddings for mouse intestinal cells ^21^ without pathogen treatment (candidate background dataset), colored by cell type. Middle: UMAP coordinates of scVI-derived embeddings for mouse intestinal cells ^21^ exposed to pathogen treatment, colored by cell type (target dataset). Right: Middle panel colored by the pathogen type. **b**. Left: Venn diagram illustrating the designed relationship between target and candidate background datasets, along with the corresponding BasCoD p-value and the decision regarding candidate background validity. Middle: UMAP embeddings showing contrastiveVI salient features of the target dataset contrasted against the candidate background, colored by cell type. Right: Same embedding as the middle panel, colored by the pathogen type.

**Supplementary Figure 4.**
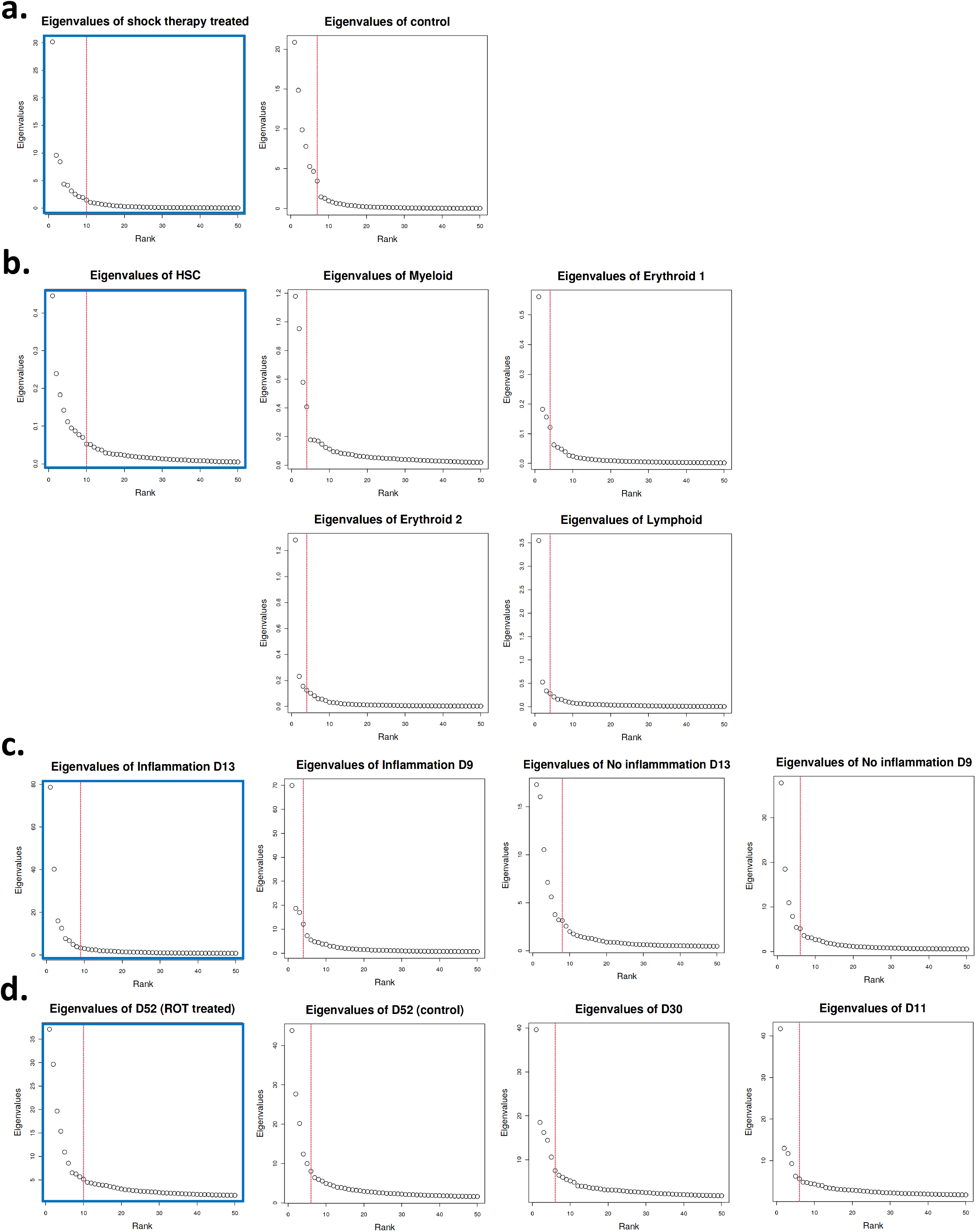
**a-d**. Eigenvalue scree plots of the target dataset and candidate background datasets for mouse protein^10, 11^ (**a**), HCA-BM^18^ (**b**), iPSCs^1^ (**c**)and HSPCs datasets (**d**). The scree plot of the target dataset is boxed in as blue. For each scree plot, the red vertical line depicts the rank selected for the dataset.

**Supplementary Figure 5.**
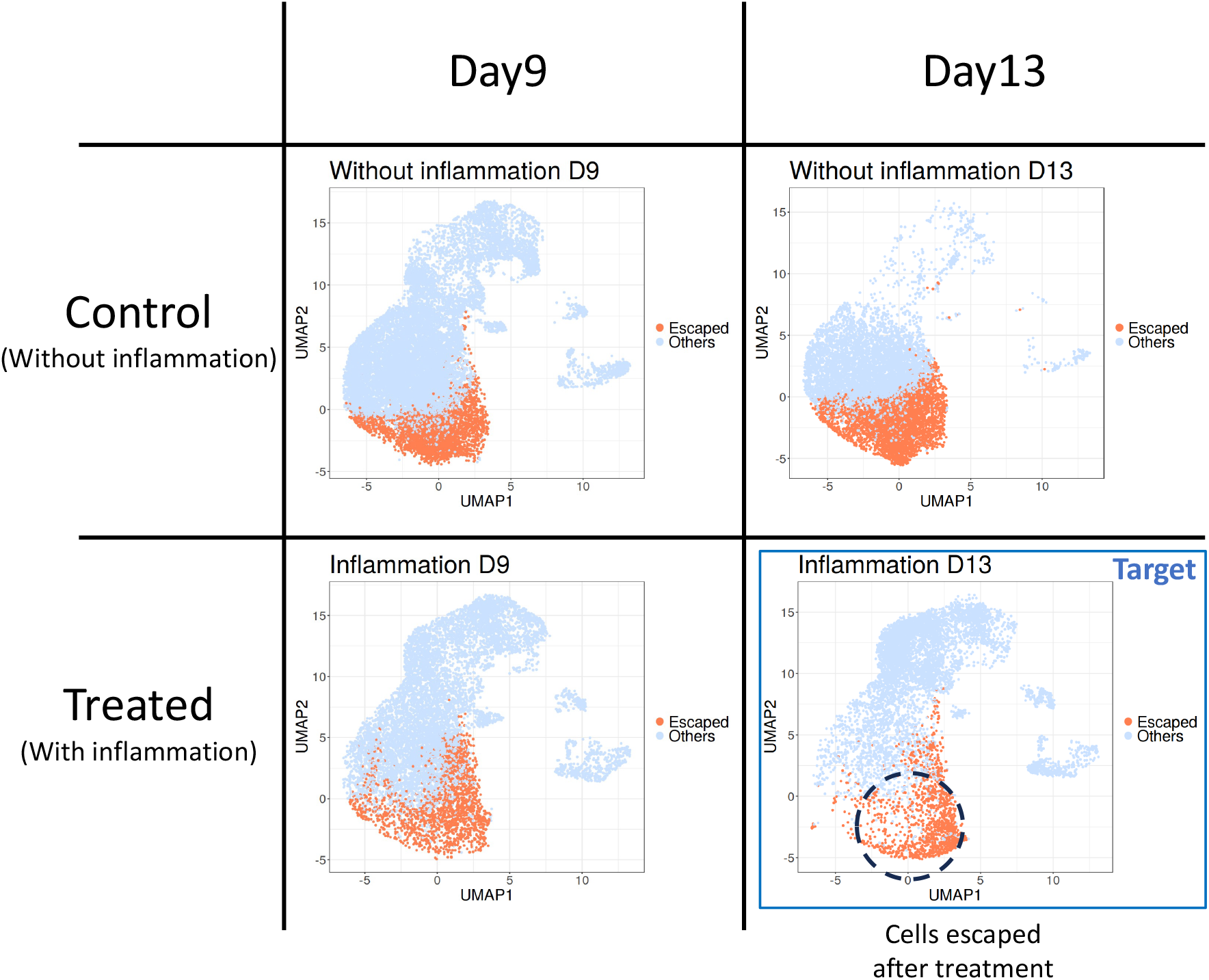
UMAP coordinates of the HSPC cells under conditions {D9 without inflammation, D9 with inflammation, D13 without inflammation, D13 with inflammation}. D13 cells with inflammation (blue boxed panel) was defined as target dataset. Cells that differentiated by D13 after inflammation (orange colored cells within the dashed-lined circle) were defined as “escaped cells”.

**Supplementary Figure 6.**
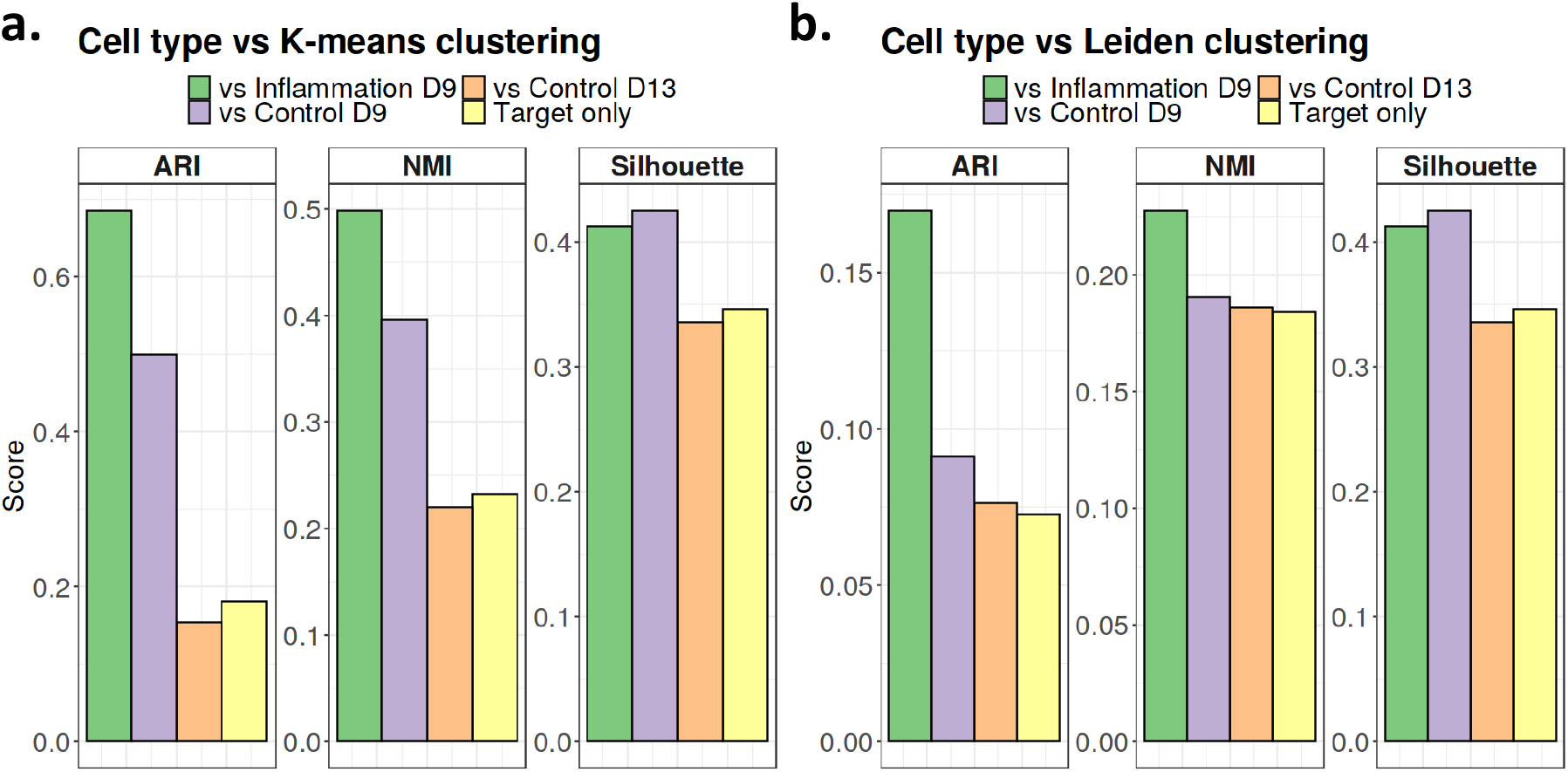
**a**,**b**. Numerical evaluation of the K-means (or Leiden) clustering results for each cPCA conducted for HSPC cells in **Fig**. 5a. ARI, NMI, and average Silhouette scores were used for the evaluations in each panel.

**Supplementary Figure 7.**
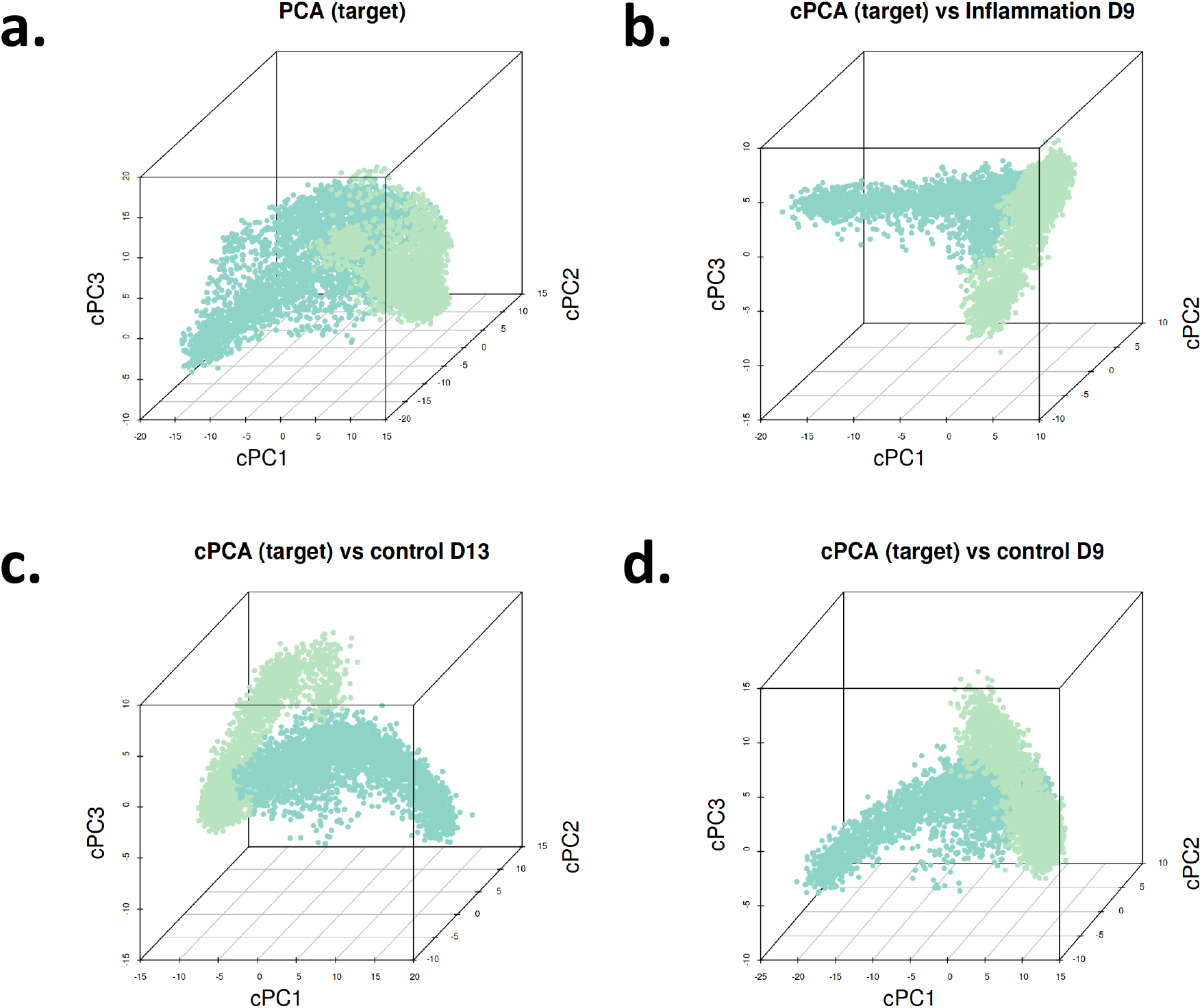
Leiden clustering result on the PCs of the target dataset of HSPC cells (**a**), and cPCs of the target dataset against each contrast (candidate background dataset) (**b-d**).

**Supplementary Figure 8.**
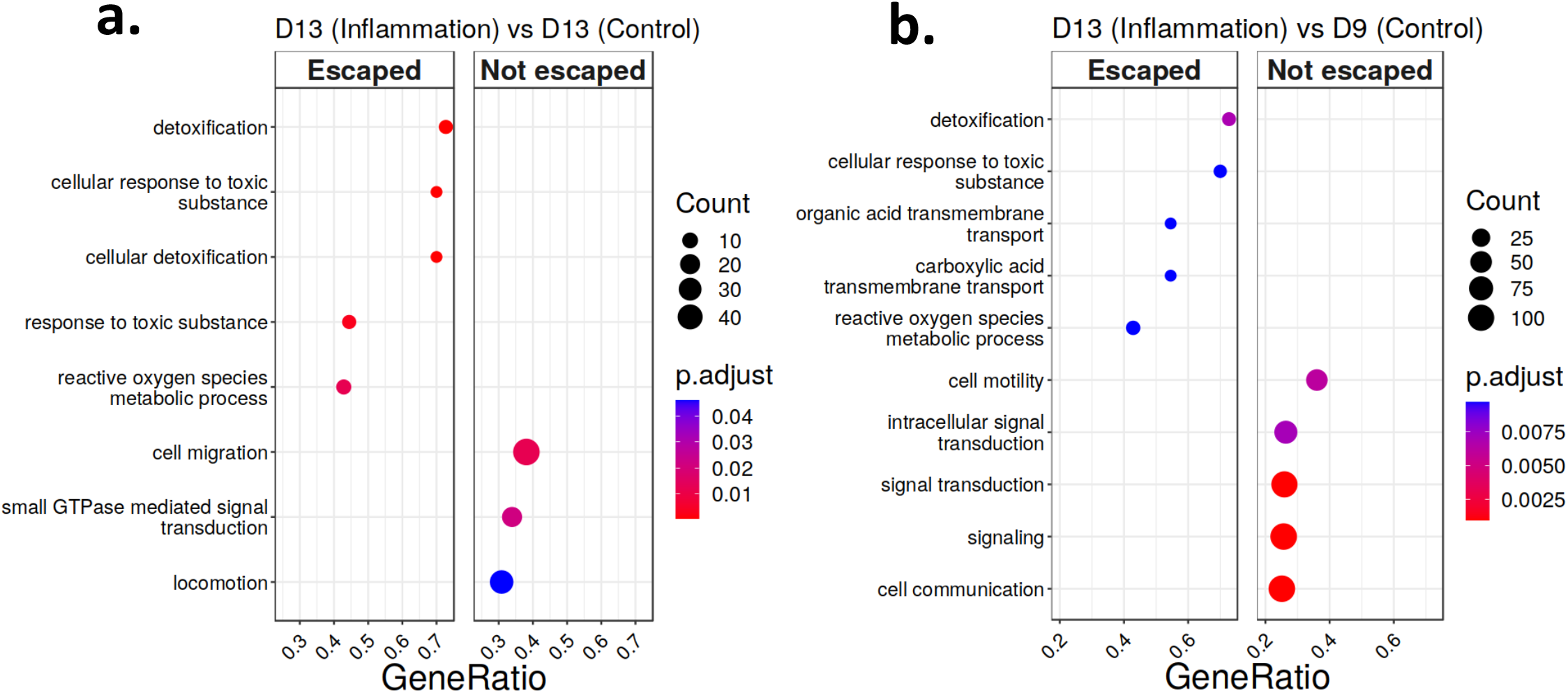
**a**. Gene set Enrichment Analysis (GSEA) comparing escaped and not escaped cells of the target dataset (D13 with inflammation). The “Escaped” panel contains the gene functions enriched in escaped cells and the “Not escaped” panel contains that enriched in not escaping cells. **b**,**c**. Gene set enrichment analysis comparing the target dataset’s two cell clusters derived from contrastive PCA embeddings when the target dataset is contrasted against inflammation-treated D13 cells (or D9 without inflammation).

## Notes

### Competing Interest Statement

The authors have declared no competing interest.

### Funding

- National Institutes of Health: HG003747, HG012881
- Chan Zuckerberg Initiative

## References

1. Jerber, J., Seaton, D.D., Cuomo, A.S., Kumasaka, N., Haldane, J., Steer, J., Patel, M., Pearce, D., Andersson, M., Bonder, M.J., et al.: Population-scale single-cell rna-seq profiling across dopaminergic neuron differentiation. Nature genetics 53(3) (2021) 304–312

2. Soskic, B., Cano-Gamez, E., Smyth, D.J., Ambridge, K., Ke, Z., Matte, J.C., Bossini-Castillo, L., Kaplanis, J., Ramirez-Navarro, L., Lorenc, A., et al.: Immune disease risk variants regulate gene expression dynamics during cd4+ t cell activation. Nature genetics 54(6) (2022) 817–826

3. Dixit, A., Parnas, O., Li, B., Chen, J., Fulco, C.P., Jerby-Arnon, L., Marjanovic, N.D., Dionne, D., Burks, T., Raychowdhury, R., et al.: Perturb-seq: dissecting molecular circuits with scalable single-cell rna profiling of pooled genetic screens. cell 167(7) (2016) 1853– 1866

4. Abid, A., Zhang, M.J., Bagaria, V.K., Zou, J.: Exploring patterns enriched in a dataset with contrastive principal component analysis. Nature communications 9(1) (2018) 2134

5. Abid, A., Zou, J.: Contrastive variational autoencoder enhances salient features. arXiv preprint arXiv:1902.04601 (2019)

6. Severson, K.A., Ghosh, S., Ng, K.: Unsupervised learning with contrastive latent variable models. In: Proceedings of the AAAI Conference on Artificial Intelligence. Volume 33. (2019) 4862–4869

7. Weinberger, E., Lin, C., Lee, S.I.: Isolating salient variations of interest in single-cell data with contrastivevi. Nature Methods 20(9) (2023) 1336–1345

8. Weinberger, E., Covert, I., Lee, S.I.: Feature selection in the contrastive analysis setting. Advances in Neural Information Processing Systems 36 (2024)

9. Zhang, B., Nyquist, S., Jones, A., Engelhardt, B.E., Li, D.: Contrastive linear regression. arXiv preprint arXiv:2401.03106 (2024)

10. Ahmed, M.M., Dhanasekaran, A.R., Block, A., Tong, S., Costa, A.C., Stasko, M., Gardiner, K.J.: Protein dynamics associated with failed and rescued learning in the ts65dn mouse model of down syndrome. PloS one 10(3) (2015) e0119491

11. Higuera, C., Gardiner, K.J., Cios, K.J.: Self-organizing feature maps identify proteins critical to learning in a mouse model of down syndrome. PloS one 10(6) (2015) e0129126

12. Norman, T.M., Horlbeck, M.A., Replogle, J.M., Ge, A.Y., Xu, A., Jost, M., Gilbert, L.A., Weissman, J.S.: Exploring genetic interaction manifolds constructed from rich single-cell phenotypes. Science 365(6455) (2019) 786–793

13. Lopez, R., Regier, J., Cole, M.B., Jordan, M.I., Yosef, N.: Deep generative modeling for single-cell transcriptomics. Nature methods 15(12) (2018) 1053–1058

14. Wilson, D.J.: The harmonic mean p-value for combining dependent tests. Proceedings of the National Academy of Sciences 116(4) (2019) 1195–1200

15. Trapnell, C., Cacchiarelli, D., Grimsby, J., Pokharel, P., Li, S., Morse, M., Lennon, N.J., Livak, K.J., Mikkelsen, T.S., Rinn, J.L.: Pseudo-temporal ordering of individual cells reveals dynamics and regulators of cell fate decisions. Nature biotechnology 32(4) (2014) 381

16. Ji, Z., Ji, H.: Tscan: Pseudo-time reconstruction and evaluation in single-cell rna-seq analysis. Nucleic acids research 44(13) (2016) e117–e117

17. Street, K., Risso, D., Fletcher, R.B., Das, D., Ngai, J., Yosef, N., Purdom, E., Dudoit, S.: Slingshot: cell lineage and pseudotime inference for single-cell transcriptomics. BMC genomics 19 (2018) 1–16

18. Hou, W., Ji, Z., Chen, Z., Wherry, E.J., Hicks, S.C., Ji, H.: A statistical framework for differential pseudotime analysis with multiple single-cell rna-seq samples. Nature communications 14(1) (2023) 7286

19. Yu, G., Wang, L.G., Han, Y., He, Q.Y.: clusterprofiler: an r package for comparing biological themes among gene clusters. Omics: a journal of integrative biology 16(5) (2012) 284–287

20. Hawke, S., Ma, Y., Li, D.: Contrastive dimension reduction: when and how? Advances in Neural Information Processing Systems 37 (2024) 74034–74057

21. Haber, A.L., Biton, M., Rogel, N., Herbst, R.H., Shekhar, K., Smillie, C., Burgin, G., Delorey, T.M., Howitt, M.R., Katz, Y., et al.: A single-cell survey of the small intestinal epithelium. Nature 551(7680) (2017) 333–339

22. Liu, Y., Xie, J.: Cauchy combination test: a powerful test with analytic p-value calculation under arbitrary dependency structures. Journal of the American Statistical Association 115(529) (2020) 393–402

